# Identification of specific gut bacteria and metabolomic signatures related to Myocardial Insulin Resistance in Type 2 Diabetes

**DOI:** 10.64898/2026.01.08.698405

**Authors:** Xingpeng Xiao, Andreea Ciudin, Rafael Simó, Carolina Aparicio, Martina Palomino-Schätzlein, José Raul Herance

**Author notes:** Correspondence (J.R.H.); (M.P.-S.); (ext. 4946) (J.R.H.); (M.P.-S.). (X.X.); (C.A.) (A.C.); (R.S.).

## Abstract

**Background:** In myocardial insulin resistance (mIR), insulin fails to appropriately stimulate myocardial glucose uptake, which can contribute to the development of heart disease. This reversible metabolic abnormality occurs in only a fraction of patients with type 2 diabetes mellitus (T2DM) and may be present even before clinical symptoms or the onset of diabetes. Identifying biomarkers of early-stage mIR is crucial to halt its progression with appropriate treatment and lifestyle interventions. In this context, we explore, for the first time, specific gut bacteria and fecal and plasma metabolic signatures associated with mIR.

**Methods:** Forty-two T2DM patients were divided into two groups, mIR patients patients with myocardial insulin sensitivity (mIS), depending on their myocardial glucose uptake measured with ^18^F-FDG PET during the hyperglycemic clamp. For each group, fecal microbiota was quantified by Illumina sequencing, and metabolomic profiles in feces and plasma were determined by NMR spectroscopy.

**Results:** The Faith’s phylogenetic diversity index was reduced significantly in mIR patients, compared with mIS patients. Linear discriminant analysis effect size (LEfSe) analysis revealed 25 bacterial taxa differing between mIS and mIR groups, and 6 bacteria (*Thermicanus, Lachnobacterium, Desulfovibrio, Desulfuromusa, Desulfosporosinus* and *Coprobacillus cateniformis*) were significantly decreased in mIR patients at the genus and species levels. Multivariate metabolomic analysis determined that six fecal metabolites (arabinose, β-galactose and ribose, aspartate, histidine and lactate) and six plasma metabolites (creatinine, myo-inositol, threonine, β-galactose, glycerol and choline) were reduced in mIR patients while plasmatic levels of VLDL(CH_3_) were elevated significantly. Myocardial IR correlated positively with plasmatic VLDL(CH_3_) and was negatively associated with two bacteria (*Desulfosporosinus* and *Coprobacillus cateniformis*), five fecal metabolites (arabinose, β-galactose, ribose, aspartate, and histidine) and two plasma metabolites (choline and β-galactose).

**Conclusions:** Our pilot study proved that the gut microbiome and the metabolic plasma profile is altered in T2DM patients with mIR. Regression analysis and predictive modelling identified the bacteria *Desulfosporosinus* and the metabolite galactose as key biomarkers, which could aid in the identification of T2DM patients at risk for myocardial IR and in the design of novel therapeutic strategies to prevent it.

## 1 Introduction

Diabetes is a global public health problem that severely affects both patients and healthcare systems. The International Diabetes Federation estimated that in 2024, 589 million people (aged 20–79) had diabetes, representing 1 in 9 adults, and an alarming increase to 853 million is projected by 2035, 1 in 8 adults [1]. Over 90% of people with diabetes have type 2 diabetes mellitus (T2DM), caused mainly by bad lifestyle habits. T2DM is featured due to the relative insulin deficit caused by pancreatic β-cell dysfunction and/or by the loss of insulin sensitivity (IS) to incorporate glucose by cells, insulin resistance (IR), in key target organs or tissues such as heart and skeletal muscle. [2]. IR usually triggers several pathological processes in these organs and tissues due to the activation of new pathways for cell survival [3] as well as the increase of blood glucose levels, inducing hyperglycemia, and causing micro- and macrovascular pathological effects that can even cause death. The cardiovascular (CV) effects of T2DM are the main cause of death in this type of patients and is closely related with IR in the myocardium [4]. Therefore, it is mandatory to delve deeper into the causes of this IR-CV complications promoted by T2DM and find biomarkers and therapeutic approaches for precision medicine of these patients.

Myocardium, the striated muscle of the heart, is responsible for its contractile function, requiring a constant supply of energy. Its main source of energy is fatty acids, but it also requires glucose for its appropriate functioning [5]. Myocardial IR may lead to post-ischemic heart failure or other complications, being an independent risk factor for heart failure [6, 7]. Despite this, not all T2DM patients with systemic IR suffer from myocardial IR. A recent study from our group showed that two phenotypes of T2DM patients coexisted, myocardial IR (mIR) and myocardial IS (mIS), being around the 60% and 40% of all T2DM patients, respectively [8]. Moreover, the systemic homeostasis model assessment of insulin resistance (HOMA-IR), commonly used for IR assessment in T2DM, works exclusively for the mIR phenotype, which features high coronary artery calcification (CAC) scores and, therefore, higher cardiovascular risk [8]. On the other hand, to learn more about the causes and connections of myocardial IR with diabetic cardiomyopathy and related complications, studies using both phenotypes are mandatory. Among all interesting studies, the relationship between the gut microbiota (GM) and its metabolites with the specific myocardial IR are one of the most promising ones for its possible clinical consequences. Therefore, these will be the goals of the current study.

GM is an assortment of microorganisms including bacteria, archaea, fungi, viruses and bacteriophages, inhabiting the gastrointestinal tract of mammals and forming a dynamic and spatially heterogeneous ecosystem [9]. The development of sequencing technology and bioinformatics analysis technology has deepened the GM understanding, especially after the launch of the Human Microbiome Project and MetaHit [10, 11]. Plenty of studies have proven that GM plays many crucial roles for our health, including participating in the digestive process, strengthening the gut barrier, protecting against pathogens and regulating host immunity [12]. Hence, to some extent, the GM is considered as an “organ” of human body [9]. Nowadays, there is strong evidence that GM is closely related with the development of many diseases, including obesity and T2DM. Patients with T2DM were found to have a moderate degree of GM misbalance, manifesting a reduction of some butyrate-producing bacteria and an increase of some opportunistic pathogens such as *Escherichia coli* [13]. Moreover, these butyrate-producing bacteria were linked with IR, as multiple studies revealed that butyrate may improve IR and increase energy expenditure in obese patients as well as in T2DM [14, 15]. In addition, other T2DM studies also demonstrated that GM and its related metabolites were associated with systemic IR [16, 17]. On the other hand, endotoxin producing bacterium in the gut of obese mice such as *Escherichia coli* could induce obesity and IR through continuous subcutaneous infusion of lipopolysaccharide [18], as well as GMs related with the production of short-chain fatty acids such as *Eubacterium ventriosum* and *Roseburia intestinalis* in obesity [19] and *Faecalibacterium prausnitzii*, *Roseburia intestinalis* and *Eubacterium rectale* in T2DM [13].

At another level, metabolomic analysis provides a valuable complement to GM sequencing by offering functional insights into microbial activity, capturing in part the biochemical output of microbial metabolism. It enables the identification and quantification of small-molecule metabolites produced by different microbial strains, host–microbe interactions, and dietary influences, that play key roles in host physiology and disease states [20, 21]. Thus, by integrating sequencing with metabolomic profiling, we can gain a more comprehensive understanding of the functional state and dynamics of the gut microbiome [22, 23]. This picture can further be complemented with a non-invasive blood plasma metabolomics analysis, that can reflect the general state of the organism and is less influenced by dietary conditions. Previous studies had unveiled that alterations of some metabolites in serum or feces were associated with T2DM and IR. For instance, high levels of branched-chain amino acids in plasma were related with an increased risk of T2DM and IR [24]. In feces, T2DM patients had reduced propionic acid, valeric acid and butyric acid as well as increased succinate [25]. However, to our knowledge, no study has focused on gut microbiome genomics or metabolomics specifically targeting myocardial insulin resistance, despite the fact that its consequences are well known, that it is not fully aligned with systemic insulin resistance indices, and that distinct myocardial phenotypes have been described [8, 26]. Therefore, this is a challenge that will be the objective of our study in order to clarify the effect of GM and related metabolites on myocardial IR in T2DM patients.

## 2 Materials and Methods

### 2.1 Study Design

Forty-seven Caucasian patients diagnosed with T2DM and regular consumers of the Mediterranean diet were recruited at the Endocrinology Department of Vall d’Hebron University Hospital.

The inclusion criteria were: (1) at least 5 years of T2DM diagnosis, (2) more than 1 year of T2DM controlled and (3) age between 47-79 years. The exclusion criteria were: (1) type 1 diabetes, (2) any previous CV event, (3) contraindication for PET/CT, (4) smokers or ex-smokers who have stopped smoking for less than 1 year, (5) regular alcohol drinker (>40 gr/day), and (6) any pathology related to a short life expectancy.

This study was conducted according to the tenets of the Helsinki Declaration after approval by the Ethic Committee of the Vall d’Hebron University Hospital (ClinicalTrials.gov: NCT02248311). Written informed consent of all participants was collected.

### 2.2 Myocardial insulin-sensitivity quantification

This assessment has been performed following a methodology previously described by our group [8]. Briefly, two ^18^F-FDG PET/CT scans were performed for each patient, at baseline and after a hyperinsulinemic-euglucemic clamp (HEC). These scans were performed in a random order within 2 days under at least 8 h of fasting conditions and 24 h withdrawal of medication. After PET/CT acquisition, myocardial IS, the inverse of IR, were determined by quantifying the difference in standardized uptake value (ΔSUV) between both PET/CT scans (SUV_HEC_ − SUV_BASELINE_) [26]. Thus, two phenotypes according to the uptake of ^18^F-FDG in the myocardium were determined [8, 26]. Twenty-one patients exhibited a clear enhancement of ^18^F-FDG uptake and twenty-six patients showed no or a marginal increase after the HEC procedure, classifying patients into the myocardial insulin-sensitivity (mIS) and myocardial insulin-resistance (mIR) phenotypes, respectively [8, 26]. Finally, due to the lack of fecal samples, only forty-two patients were included in the study: seventeen patients with mIS and twenty-five patients with mIR.

### 2.3 Anthropometric and Biochemical Measurements

Anthropometric measurements were conducted for all participants at the Nuclear Medicine Department, including body weight, height, waist and hip circumferences before the PET/CT scan. Blood samples were collected under fasting conditions and analyzed at the Biochemistry Core Facilities of Vall d’Hebron University Hospital using standardized and validated routine protocols. The following parameters were determined: plasma glucose, glycated haemoglobin (HbA1c), high-density lipoprotein (HDL), low-density lipoprotein (LDL), triglycerides (TG), total cholesterol, C-peptide, total free fatty acids, fructosamine, adiponectin, sodium, potassium, phosphate, calcium, chloride, total protein, albumin and creatinine. Non-routine parameters such as insulin, leptin, interleukin-6 (IL-6), troponin I were also measured following standardized methodologies previously described [27]. The HOMA-IR and the triglyceride-glucose (TyG) index were calculated by using the described formula [28, 29]. Systemic insulin sensitivity (IS_HEC_) was measured by the mean glucose infusion rate during the last 40 minutes of the HEC procedure [30, 31].

### 2.4 Sample Preparation for Metabolomics

Blood samples were collected from patients under fasting conditions and stored at 4 °C. They were processed within 1 h by following the protocol described previously by our group [32]. Briefly, 2 mL of Histopaque^®^ 1119 and 1077 (Sigma-Aldrich, UK) were successively added to a 15-mL clean centrifuge tube. Then, blood samples were added after gentle shaking and then centrifuged at 300 g and 4 °C for 30 min. The upper light-yellow layer (plasma) was transferred into a 2 mL clean Eppendorf tube and stored at −80 °C until its metabolomic analysis.

Stool samples were collected once the patients arrived at the Nuclear Medicine Department, aliquoted and stored at −80 °C immediately. For the extraction of fecal metabolites, a previous described protocol was followed [33]. Briefly, frozen fecal samples were thawed on ice, and 100 mg feces were weighed in a 1.5 mL plastic tube and then homogenized with 1 mL 0.1M phosphate D_2_O buffer (with 5 mM TSP, pH 7.4). Samples went through 3 freeze-and-thaw cycles by alternate immersion into liquid nitrogen and tap water, and subsequently were centrifuged at 15000 g for 15 min after a 15-second vortex. The supernatant was collected and stored at −80 °C until analysis.

### 2.5 NMR Spectroscopy

Sample preparation, NMR acquisition, and analysis were carried out following our established protocols [33, 34]. In brief, plasma samples were thawed on ice before NMR analysis. Then, 300 μL of plasma was combined with 300 μL of 10% D_2_O buffer containing 5 mM TSP, 140 mM phosphate, and 0.04% NaN_3_, at pH 7.4. Subsequently, 550 μL of the resultant mixture was transferred into a 5-mm NMR tube for its analysis. The fecal supernatant, was thawed on ice, and 600 μL were directly transferred into the NMR tube. ^1^H-NMR spectra were acquired at 27 °C using a Bruker 600 MHz spectrometer (Rheinstetten, Germany) equipped with a TCI cryoprobe and processed with TopSpin 3.2 software (Bruker). D_2_O served as the solvent of the field frequency, a 4 s relaxation delay was implemented between free induction decays (FIDs), and water suppression by presaturation was applied. Digitalization was performed over 64 K data points across a spectral width of 30 ppm for optimal baseline correction. To quantify lipoproteins, additional diffusion-edited experiments were conducted to minimize interference from small metabolite signals. FID values were subjected to exponential function multiplication with a 0.5 Hz line broadening factor. The reference for plasma and feces were the α-glucose anomeric doublet (δ 5.23) and the TSP signal (0), respectively. ^1^H-NMR signals were assigned to their corresponding metabolites using 2D NMR experiments and spectral databases such as the Human Metabolome Database (HMBD) [35] and Biological Magnetic Resonance Data Bank (BMRB) [36]. In cases of ambiguity, assignments were validated by adding a reference compound to the sample and repeating spectra to confirm signal matches. Normalization of all spectra to total intensity was conducted to minimize concentration differences and experimental errors. Integration was carried out using Global Spectral Deconvolution in MestreNova 8.1 (Mestrelab Research, Spain), with optimal integration regions defined for each metabolite.

### 2.6 Extraction of Fecal DNA

The procedure for fecal DNA extraction was adapted from Yu *et al* [37]. Briefly, frozen fecal samples were scraped and weighed 300∼500 mg. Immediately, 1 mL lysis buffer (500 mM NaCl, 50 mM Tris-HCl, pH 8.0, 50 mM EDTA, and 4 % sodium dodecyl sulfate) was added to a 2 mL Eppendorf tube containing the feces. Samples were homogenized by resuspension using a micropipette and vortexed, then heated at 88 °C for 15 min, with gentle shaking every 5 min. The supernatant was harvested by centrifugation at 16000 g and 4 °C for 5 min. Then, the precipitate was resuspended with 300 μL of fresh lysis buffer, and then the above incubation and centrifugation steps were repeated. Both supernatants were pooled together, and steps 6 to 10 of the extraction protocol of Yu *et al* were applied. Then, fecal DNA dissolved in Tris-EDTA buffer was purified by QIAamp® Fast DNA Stool Mini Kit (Catalog no. 51604, Qiagen, Germany). Briefly, the above dissolved fecal DNA solution (about 200 μL) was added to 700 μL InhibitEX Buffer and vortexed for 1 min. Then, the mixed solution was incubated on a heating block at 70 °C for 5 min, vortexed for 15 s, and centrifuged at 10000 g for 2 min. The supernatant was divided into four 1.5 mL Eppendorf tubes with a volume of 200 μL. Each tube was added 1.5 μL DNase-free RNase (10 mg/mL) and incubated at 37 °C for 15 min. Subsequently, 7.5 μL of proteinase K and 200 μL Buffer AL were added and then the protocol “Isolation of DNA from Stool for Pathogen Detection”of the above Qiagen kit was followed from step 7 to the end. Solutions from the same stool passed sequentially through the QIAamp spin columns before adding Buffer AW1, both from the kit.

### 2.7 Illumina Sequencing

Before applying sequencing, the quality and concentration of extracted genomic fecal DNA were measured. The purity and concentration of fecal DNA was determined by Nanodrop 2000 spectrophotometer (ThermoFisher Scientific, USA), and a 260/280 ratio of 1.96 ± 0.07 (mean ± SD) was considered pure. Moreover, the extracted DNA was checked by agarose gel electrophoresis and PCR with 16S universal primers respectively.

Fecal DNA samples were sequenced in ATLAS Biolabs GmbH (Berlin, Germany) using their Illumina MiSeq platform using a standardized protocol [38] targeting the V4 region of the 16S rRNA gene. Each sample was amplified in triplicate and the amplification products were pooled together. The expected size of amplicons was around 300–350 bp. The amplicons of all samples were verified by agarose gel and quantified by Quant-iT PicoGreen dsDNA assay (Invitrogen, USA). The 16S library was set up by combining an equal amount of amplicon from each sample (240 ng) and cleaned by using UltraClean PCR Clean-Up Kit (Mo Bio, USA). Then, one aliquot of the library was sequenced on the MiSeq platform with 5-10% PhiX.

Raw sequence reads were processed according to the BaseSpace 16S Metagenomics App (Illumina) and QIIME 2 pipelines [39]. Fastq documents were annotated by the rdp_16s_v18 database on VSEARCH (v2.21.1) to generate an operational taxonomic unit (OTU) table at a similarity threshold of 97% [40]. The raw sequence data for the 16S rRNA gene can be accessed from NCBI Sequence Read Archive (SRA) database with accession ID PRJNA810955.

### 2.8 Data Analysis

For sequencing data, using the OTU table, Venn diagram and principal coordinate analysis (PCoA) were made to see the structural differences of microbial community of fecal samples by two platforms, Wekemo Bioincloud [41] and ImageGP [42]. Using data on the relative abundance of microorganisms detected in most fecal samples, linear discriminant analysis effect size (LEfSe) was performed on ImageGP to discover potential biomarkers. For metabolomic data, multivariate analysis such as orthogonal-orthogonal projection to latent structure discriminant analysis (OPLS-DA) or partial least squares discriminant analysis (PLS-DA) were performed to reduce the dimensionalities and identify the most important metabolites by using SIMCA-P software (version 14.1) (Umetrics, Sweden). OPLS-DA and PLS-DA models were validated by the permutation plot (100 folds) and the reliability of these models was assessed by ANalysis Of VAriance testing of Cross-Validated predictive residuals (CV-ANOVA) in SIMCA-P. During the OPLS-DA or PLS-DA analysis, variables with VIP values > 1.0 were selected for further statistics.

After specific gut bacteria and metabolites were determined, their relationships were examined in all patients as well as in patients with a single phenotype by network analysis using Spearman’s rank correlation. Network analysis was carried out on the Wekemo Bioincloud platform [41]. Statistical analysis was performed in GraphPad Prism 8.0.1 (San Diego, USA). If the data follow a normal distribution, they were expressed as mean ± standard error of the mean (SEM), otherwise, they were shown as median (interquartile range). Differences with p<0.05 were considered significantly different between groups, and for correlations, a p<0.05 was considered a significant correlation between variables.

## 3 Results

First, a comparison of the anthropometric and biochemical data was conducted between the two T2DM phenotypes, mIS and mIR, that were age and sex matched (Table 1). No significant differences in anthropometric parameters such as the waist/hip ratio and body mass index (BMI) between the two groups were found. As expected, IS_HEC_ was significantly lower in mIR patients (p=0.004), and therefore HOMA-IR tended to increase in mIR (p=0.089). Although concentrations of HDL-Cholesterol and total cholesterol did not show significant differences, the ratio of them was higher in mIR (p=0.043). Moreover, we also detected prominent differences in total proteins, albumin and electrolytes such as calcium and chloride (p<0.05) between both phenotypes.

**Table 1.** Main anthropometric and biochemical parameters of T2DM patients with mIS and mIR phenotypes.

| Parameter | mIS | mIR | P value |
| --- | --- | --- | --- |
| Number of patients | 17 | 25 |  |
| Female/Male | 9/8 | 12/13 |  |
| Age | 67.12 ± 2.03 | 65.08 ± 1.28 | 0.199 |
| Waist/hip ratio | 0.98 (0.95-1.06) | 0.97 (0.92-1.04) | 0.778 |
| BMI (kg/m<sup>2</sup>) | 31.62 ± 1.12 | 31.59 ± 0.70 | 0.805 |
| HbA1c (%) | 6.96 ± 0.15 | 7.30 ± 0.18 | 0.229 |
| C-peptide (ng/mL) | 1.81 ± 0.28 | 2.03 ± 0.21 | 0.400 |
| Glucose (mg/dL) | 122.41 ± 7.96 | 130.40 ± 6.73 | 0.590 |
| Insulin (mU/L) | 15.12 (9.31-19.24) | 17.84 (11.51-33.85) | 0.279 |
| HOMA-IR | 3.88 (3.19-5.28) | 5.67 (3.68-10.36) | 0.089 |
| IS<sub>HEC</sub> [mg/(kg·min)] | 1.84 ± 0.15 | 1.34 ± 0.09 | **0.004** |
| Total cholesterol (mg/dL) | 161.2 ± 6.3 | 179.6 ± 9.0 | 0.134 |
| HDL-Cholesterol (mg/dL) | 47.9 ± 2.8 | 44.5 ± 2.1 | 0.337 |
| LDL-Cholesterol (mg/dL) | 90.5 ± 4.9 | 102.2 ± 6.9 | 0.214 |
| Triglyceride (mg/dL) | 109.0 (66.0-171.0) | 122.0 (98.5-198.5) | 0.111 |
| Total cholesterol/HDL | 3.50 ± 0.20 | 4.14 ± 0.22 | **0.043** |
| TyG index | 8.72 ± 0.14 | 9.03 ± 0.10 | 0.069 |
| Total free fatty acids (mmol/L) | 0.65 ± 0.03 | 0.76 ± 0.06 | 0.177 |
| Fructosamine (μmol/L) | 241.5 (216.0-264.0) | 258.0 (246.5-291.5) | 0.101 |
| Adiponectin (μg/mL) | 4.04 (1.93-8.41) | 3.20 (2.45-6.08) | 0.485 |
| Creatinine (mg/dL) | 0.71 (0.63-1.01) | 0.77 (0.61-0.92) | 0.854 |
| Troponin I (pg/mL) | 6.0 (3.5-10.0) | 7.0 (3.0-11.0) | 0.879 |
| IL-6 (pg/mL) | 1.87 (1.50-3.70) | 3.01 (1.81-5.89) | 0.052 |
| Total proteins (g/dL) | 6.80 (6.55-7.05) | 7.20 (6.85-7.55) | **0.013** |
| Albumin (g/dL) | 4.1 (3.9-4.3) | 4.4 (4.1-4.5) | **0.016** |
| Calcium (mg/dL) | 9.3 (9.1-9.6) | 9.7 (9.4-9.8) | **0.029** |
| Sodium (mmol/L) | 138.5 (135.8-140.1) | 138.2 (137.0-139.9) | 0.884 |
| Potassium (mmol/L) | 4.22 (4.07-4.41) | 4.17 (3.76-4.48) | 0.711 |
| Chloride (mmol/L) | 105 (103-109) | 103 (102-105) | **0.016** |

Abbreviations: BMI, body mass index; HbA1c, hemoglobin A1c; HOMA-IR, homeostasis model assessment of insulin resistance; IS<sub>HEC</sub>, insulin sensitivity measured by HEC; TyG index, triglyceride-glucose index; HDL, high-density lipoprotein; LDL, low-density lipoprotein; IL-6, interleukin-6. The P-value was calculated with unpaired t test or Mann-Whitney test. P-values < 0.05 are labelled in bold.

Then, an analysis of the genomic GM data of the patients was performed. After a standard process of GM sequencing files on VSEARCH, an OTU table was generated. In order to see the effects of mIR on GM, a Venn diagram (Figure 1A) was made. In total, 271 OTUs were detected for both phenotypes, including 59 and 18 unique to mIS and mIR respectively. Then, some alpha diversity indices were compared between groups, of which only the Faith’s phylogenetic diversity index was altered significantly and decreased in the mIR group (p<0.05). In addition, the number of observed features showed a tendency to decrease in mIR (Figure 1C). Further, to know the differences of beta diversity among samples, a PCoA plot was made but the separation of groups was only moderate (Figure 1B).

**Figure 1.**
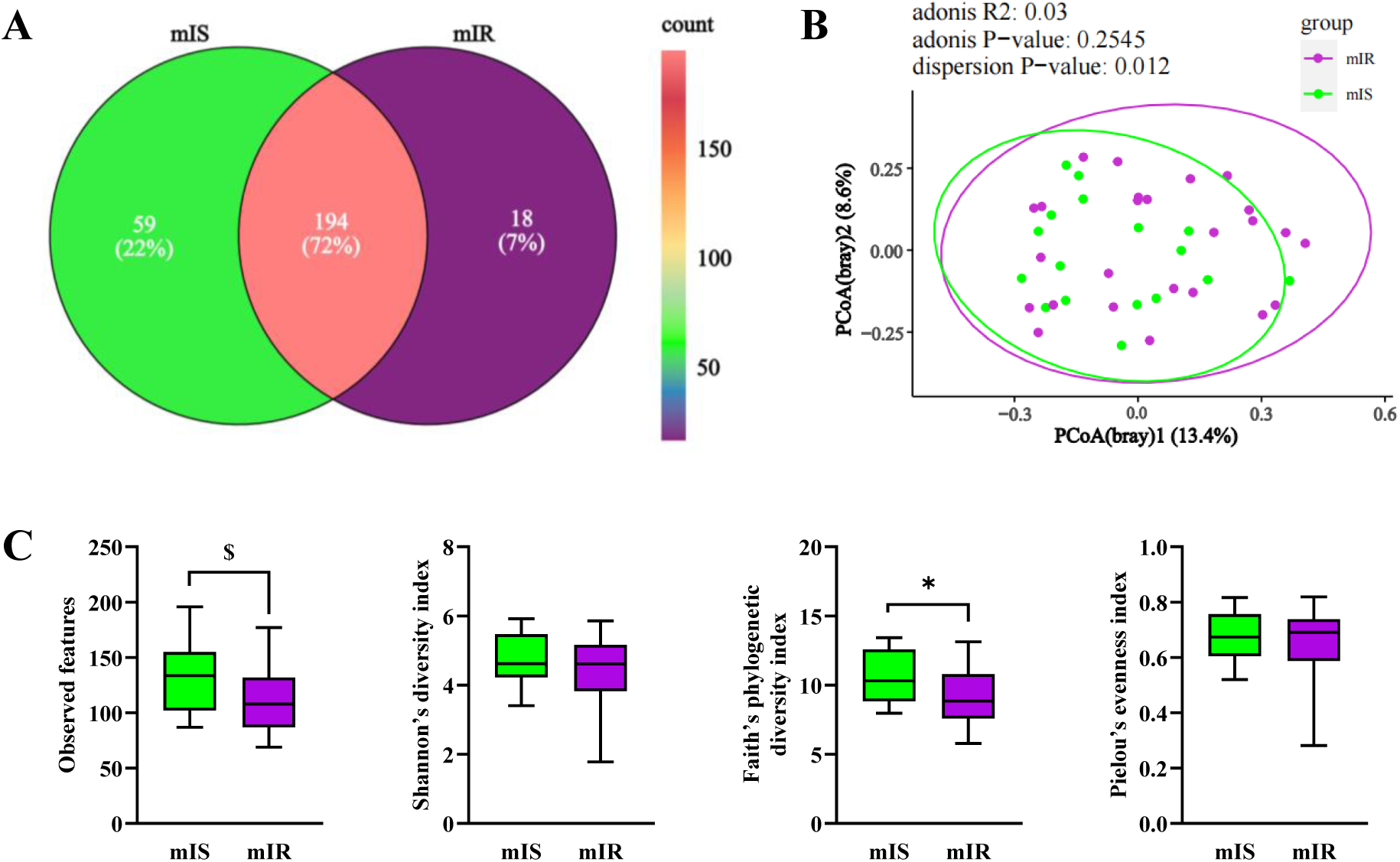
Venn diagram and biodiversity analysis. (A) Venn diagram was generated on the Wekemo Bioincloud platform by using the OTU table, (B) PCoA plot was made on the ImageGP platform by using Bray-Curtis distance method. (C) Box plots were depicted for alpha diversity indices. First three box plots reflected the species richness when the sequencing depth was 20000, and the last one presented the species evenness of a community. * and $ indicated p<0.05 and p<0.1, respectively.

Besides the biodiversity information, the relative abundance of all detected microorganisms was determined. Dominant gut bacteria were compared between both myocardial groups at phylum and family levels (Supplementary Figure 1) and showed no significant differences, only trends. Therefore, a LEfSe analysis was performed to identify any differential bacteria between groups (Figure 2A). As a result, 25 bacterial taxa were identified with a linear discriminant analysis (LDA) score >2.0, of which 20 taxa were abundant in mIS, while 5 were higher in mIR. Further, the relative abundance of these 25 taxa was statistically analyzed between two groups. At the genus and species levels, 6 bacteria (*Thermicanus*, *Lachnobacterium*, *Desulfovibrio*, *Desulfuromusa*, *Desulfosporosinus* and *Coprobacillus cateniformis*) had significant differences between mIS and mIR patients (Figure 2B) and, interestingly, all decreased in mIR (p<0.05).

**Supplementary Figure 1.**
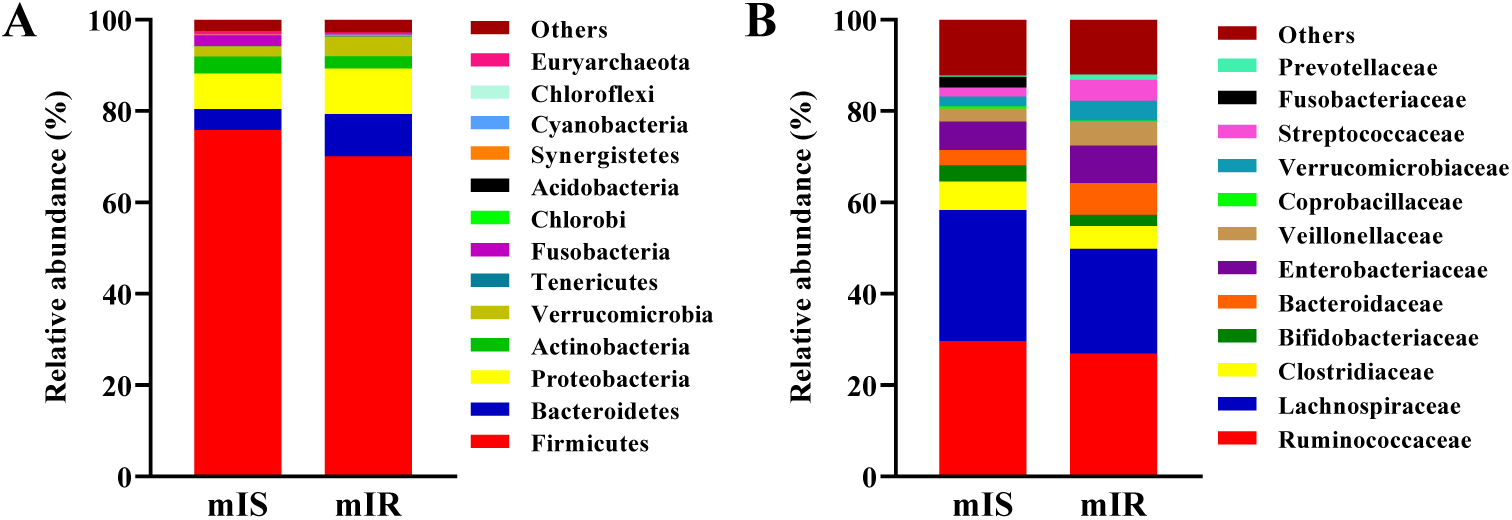
GM composition measured by Illumina sequencing. The relative abundance of dominant gut bacteria showed at the phylum level (A) and the family level (B) for both myocardial groups of patients.

**Figure 2.**
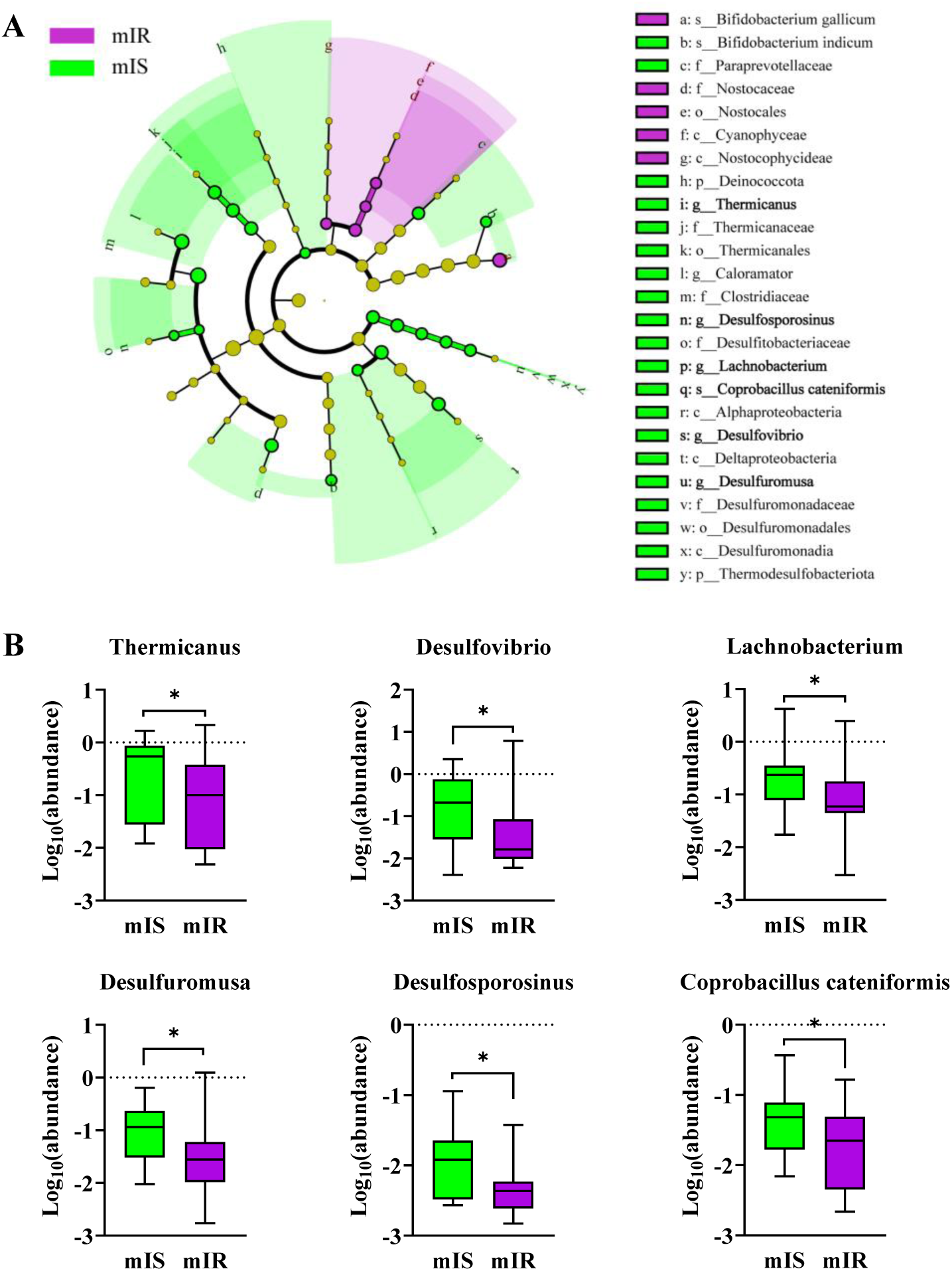
Significantly altered gut bacteria identified for patients with mIR. (A) LDA score cladogram for differentially bacterial taxa between two groups at LDA threshold >2.0. Twenty-five differentiated taxa were obtained, twenty of which were enriched in mIS while only five were higher in mIR. The letter separated by a dash before the bacterial name represented the taxonomic level. Six bacteria at lowest taxonomic levels were bold because they had significant differences between two phenotypes. (B) Box plots of significant gut bacteria at the genus and species levels. Bacterial taxa of the genus and species levels from the LDA analysis were compared, only 6 had a significant difference between groups. * indicated p<0.05.

After the identification of six GM altered between myocardial phenotypes, a metabolomic analysis of feces was performed, to identify metabolic changes that could be related with the altered bacterial strains. We first tried to build discriminant OPLS-DA or PLS-DA models between both patient groups, but no robust model could be obtained. However significant variables were identified by direct comparison of the metabolite levels between both groups. The six identified altered metabolites (Figure 3) were three monosaccharides, arabinose, β-galactose and ribose; two amino acids, aspartate and histidine; and one organic acid, lactate. All of them were significantly decreased in mIR patients (p<0.05).

**Figure 3.**
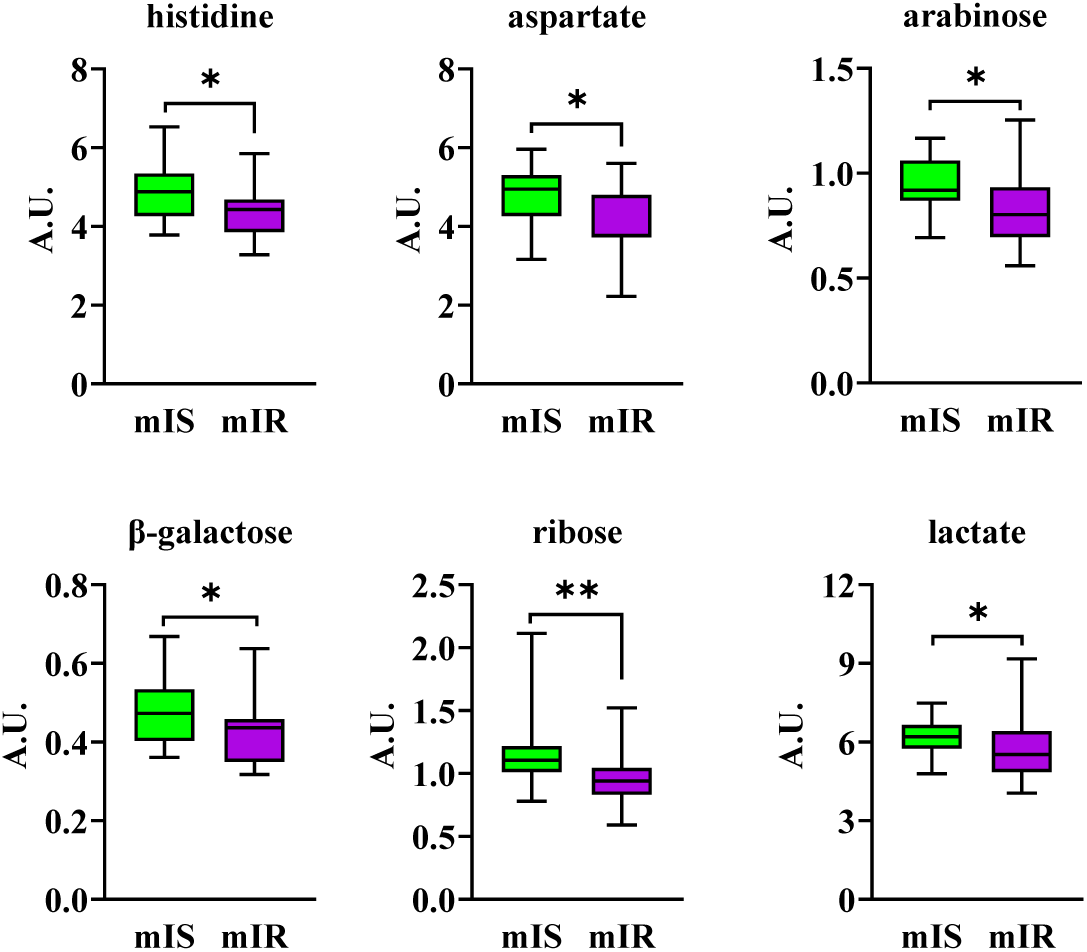
Boxplots of fecal metabolites that changed significantly for patients with the mIR phenotype. Univariate statistical analysis of normalized concentrations was conducted by using Mann-Whitney test. * and ** indicated p<0.05 and p<0.01, respectively. A.U., arbitrary units.

Furthermore, the alteration of the plasma metabolome between both T2DM myocardial phenotypes was also studied. In this case, a discriminant PLS-DA model could be obtained (Figure 4A), and the patients of both groups tended to separate in the plot. The most important altered plasma metabolites were selected by VIP value > 1 and submitted to univariate analyses. Six plasma metabolites including animo acid threonine, monosaccharide β-galactose, creatinine, myo-inositol, glycerol and choline were decreased (p<0.05) while only VLDL(CH_3_) increased (p<0.01) in mIR patients (Figure 4B).

**Figure 4.**
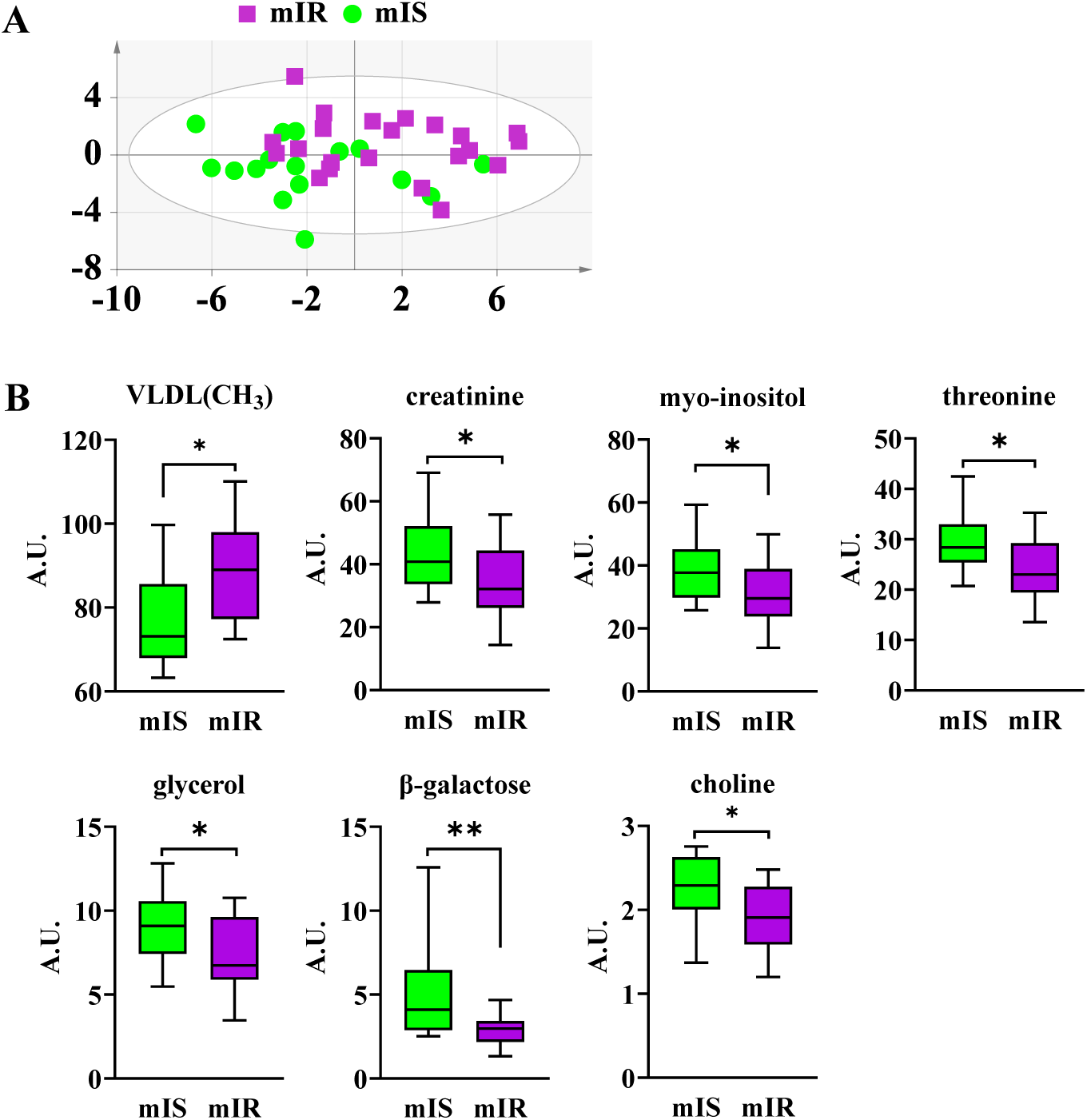
Relevant plasma metabolites for mIR patients selected by PLS-DA. (A) PLS-DA score plot of plasma metabolic profiles with model parameters: UV scaling, R2Y(cum)= 0.57, Q2(cum)= 0.218. Permutation test: R2=(0.0, 0.318), Q2=(0.0, −0.245), p from CV-Anova = 0.153. (B) Boxplot representation of significant metabolites selected by VIP value from PLS-DA (>1) and p-value from univariate analyses <0.05. * and ** indicated p<0.05 and p<0.01 respectively. A.U., arbitrary units.

After identifying the alterations of GM and the fecal and plasma metabolites between both myocardial phenotypes of T2DM patients, we assessed their associations with both systemic indices of IR and specific myocardial IR, to identify GM and metabolic correlations as well as similarities and differences of using systemic or specific parameters. For this purpose, initially Spearman’s correlation analysis was carried out using the myocardial IR of patients included in both groups (Figure 5A). For myocardial IS, positive correlations were found with the fecal metabolites arabinose, β-galactose, aspartate, histidine (p<0.05) and ribose (p<0.01); the plasma metabolites choline (p<0.05) and β-galactose (p<0.01); and the gut bacteria *Desulfosporosinus* and *Coprobacillus cateniformis* (p<0.05) as well as IS_HEC_ (p<0.01). On the other hand, myocardial IS were negatively associated with the plasma metabolite VLDL(CH_3_) (p<0.05) and HOMA-IR (p<0.01). For the systemic index of IS, IS_HEC_, some variables showed positive correlations including the fecal metabolite β-galactose and the microbial genus *Desulfosporosinus* (p<0.05). Regarding HOMA-IR, only one positive correlation with the plasma metabolite VLDL(CH_3_) (p<0.05) was shown. Of course, as expected, IS_HEC_ was always negatively associated with HOMA-IR (p<0.05). Moreover, the relationship of altered gut bacteria with fecal and plasma metabolites was determined using all patients in the study. But only one significant relationship was obtained, *Desulfuromusa* was positively associated with the fecal metabolite histidine (p<0.05). We did not find a significant correlation between fecal metabolites and plasma metabolites.

**Figure 5.**
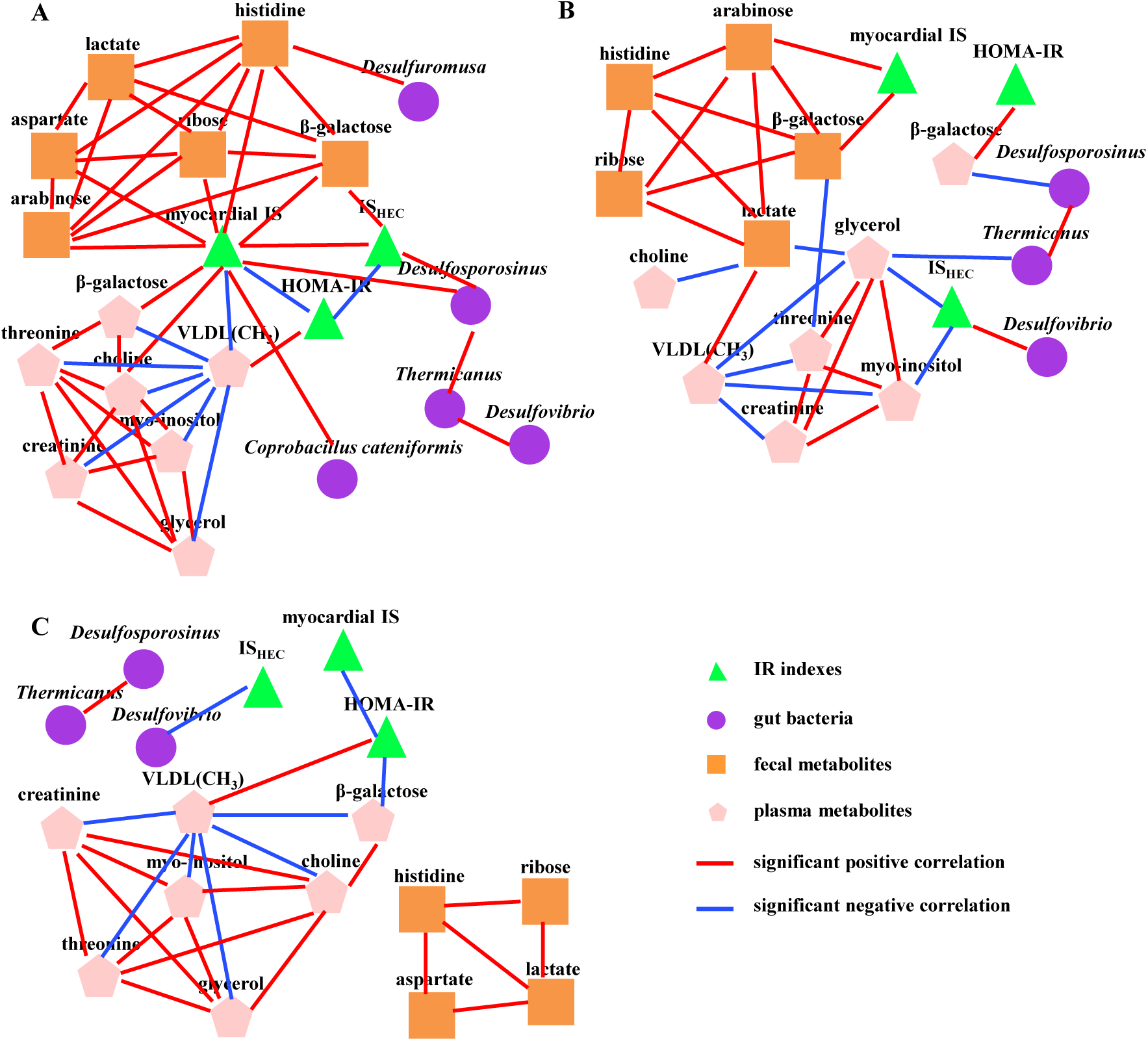
Correlation matrix of myocardial IS, IS_HEC_ and HOMA-IR with identified specific gut bacteria and metabolites. Spearman’s correlation analyses were conducted in all patients (A), in patients with the mIS phenotype (B) and with the mIR phenotype (C). Nodes with bright green, violet, orange and pink colors stand for IR indices (including myocardial IS, IS_HEC_ and HOMA-IR), gut bacteria, fecal metabolites and plasma metabolites, respectively. Red and blue lines between two nodes represented significant positive and negative correlation (p<0.05), respectively. Among identified gut bacteria and metabolites, if one did not have any significant connections with others, it was not shown in the figure.

On the other hand, we also evaluated whether these relationships still existed in patients with only mIS or mIR phenotypes. As depicted in Figure 5B and 5C, myocardial IS still had positive correlations with the fecal metabolites arabinose and β-galactose in patients with only mIS phenotype (p<0.05). HOMA-IR still showed a positive association with the plasma metabolite VLDL(CH_3_) in patients with only mIR phenotype (p<0.05).

Interestingly, several new correlations were found when analysis was performed by phenotypes separately. For instance, HOMA-IR had a positive relation with the plasma metabolite β-galactose in mIS patients, but an opposite relation in mIR patients (p<0.05). For the IS_HEC_, a positive correlation was found with *Desulfovibrio* in patients with mIS, whereas a negative correlation was observed in patients with mIR (p<0.05). Furthermore, several prominent associations were exclusively achieved in mIS patients. Thus, IS_HEC_ was negatively associated with plasma metabolites glycerol and myo-inositol (p<0.05). The fecal metabolite lactate was positively related with the plasma metabolite VLDL(CH_3_), but inversely with plasma metabolites choline and glycerol (p<0.05). The fecal β-galactose was also inversely related to the plasma threonine (p<0.05). Moreover, prominent negative correlations were displayed between the bacterium *Thermicanus* and the plasma glycerol, and between *Desulfosporosinus* and the plasma metabolite β-galactose (p<0.01).

## 4. Discussion

The current proof-of-concept clinical trial determined for the first time the differences in GM as well as plasma and fecal metabolic profiles related to myocardial IR. Thus, two coexisting myocardial phenotypes in T2DM patients, mIR and mIS, were assessed to find biomarkers and deepen the knowledge of this pathophysiological associated with CV complications [8]. In addition, the associations of specific gut bacteria with fecal and plasma metabolites were done to assess their possible relationships and to learn in more detail about their roles in myocardial IR. Finally, the associations of GM and metabolites with myocardial IR as well as systemic indices of IR were assessed to determine the usefulness of studies based on systemic indices for the specific case of the myocardium since in some cases this IR etiological entity differs from systemic to organ-specific. This is highly significant, as IR is an independent risk factor for cardiovascular mortality and heart failure.

Our results revealed clear differences between both myocardial IR phenotypes as well as between the use of myocardial IR and the systemic indices of IR. These findings highlighted the importance of a personalized management of patients with T2DM at organ level, since IR is one of the pathophysiologies that trigger important deleterious comorbidities in this type of patients [43].

### Alteration of GM in myocardial IR

The GM of T2DM patients with the mIR phenotype showed a reduction in the Faith’s phylogenetic diversity index (Figure 1). The mIR phenotype, which includes patients with a more advanced progression of T2DM, appears to differ from the mIS phenotypes that showed a more diverse GM. This result showed the importance of GM in the natural history of the evolution of myocardial and systemic IR in T2DM. To provide more details of the gut bacteria altered between mIS and mIR groups, a LEfSe analysis was used to identify their alterations between both phenotypes. Thus, six bacteria were altered at the genus and species levels. Surprisingly, all these altered bacteria were decreased in patients with the mIR phenotype (Figure 2). Regarding these bacteria, an alteration of *Thermicanus* is described here for the first time in association with T2DM patients, to our knowledge. It was only described in obese patients or centenarians, increasing its concentration against healthy controls [44, 45]. On the other hand, the alterations in butyrate-producing bacteria, such as the decrease of *Lachnobacterium* in mIR phenotype, were consistent with previous studies in T2DM [13]. Moreover, this finding is further coherent with previous works showing that a reduction in its abundance is associated with reduced IS [14].

Concerning sulfate-reducing bacteria, which can promote gut inflammation as described in T2DM, reduced gut microbiota (GM) biodiversity, and ulcerative colitis [46] the genera *Desulfovibrio* and *Desulfosporosinus* were reduced in the mIR phenotype. This finding may be consistent with previously reported alterations linking these bacteria either to changes in glucose metabolism homeostasis through hydrogen sulfide (H₂S) production [47] or, in the case of *Desulfovibrio* spp., to their enrichment in the gut of patients with T2DM [13].

Moreover, the alteration of *Desulfuromusa*, a sulfur-reducing bacterium that can reduce sulfur to H_2_S [48] has never been described in T2DM or IR, to our knowledge. However, given that it can produce H₂S, similarly to the bacteria described above, it could alter glucose homeostasis in a manner comparable to those previously described taxa. Regarding the species *Coprobacillus cateniformis*, which has not previously been associated with T2DM in patients and was reduced in the mIR phenotype, enrichment has been reported in T2DM mouse models [49], highlighting differences in gut microbiota behavior between animal models and humans, as previously proposed [50].

### Alteration of fecal metabolomics in myocardial IR

Concerning fecal metabolomics, one of the main observations is that the profile of amino acids in feces was altered, consistent with previous studies [51] that have linked alterations of amino acid metabolism in feces to T2DM and its progression [52]. In our case, the concentration of two amino acids, aspartate and histidine, was decreased in mIR patients (Figure 3). This is consistent with a longitudinal clinical trial where the aspartate deficiency was studied in the fecal analysis of 5181 Finnish men. This reduction was associated with lower insulin secretion and a higher risk of developing T2DM in humans [53]. Therefore, its depletion is associated with deficit in the insulin release and function. Our study confirms that this metabolite is also specifically related with myocardial IR in T2DM. Regarding histidine, its supplementation has been found to suppress inflammation and improve IS in obese women [54]. Therefore, this is consistent with the lower levels of this metabolite in feces of mIR patients. On the other hand, alterations of carbohydrate metabolism were also showed in the mIR patients, as proved by the reduction of the three carbohydrates β-galactose, ribose and arabinose (Figure 3). The alterations of carbohydrates and specifically these three have been described in T2DM or closed metabolic-associated diseases. For instance, D-ribose has not been described in feces of T2DM patients but in urine [55]. On the other hand, L-arabinose supplementation could alleviate IR and reduce some obesity symptoms in mice [56]. Despite that the depletion of this metabolite has not been described previously in feces of T2DM patients, to our knowledge, lower levels of β-galactose in mIR patients seems to be in concordance with previous studies showing this depletion in serum and urine [57, 58]. Lower levels of these sugars in feces in patients with the mIR phenotype suggested a change of carbohydrate metabolism linked to more advanced pathophysiology of the T2DM as the mIR phenotype.

### Alteration of plasma metabolomics in myocardial IR

Regarding the plasma metabolomics, similar to fecal metabolites, β-galactose was reduced in mIR patients. This behavior followed a similar trend in T2DM, as reported in the trials commented above. Therefore, this carbohydrate appears to play an important role in the natural history of T2DM and specifically in the development of IR. Moreover, other plasma metabolites such as myo-inositol, choline, glycerol, threonine and creatinine were also reduced and VLDL(CH_3_) increased in the mIR phenotype (Figure 4). Alterations in myo-inositol metabolism was expected since it has been associated with IR and T2DM [59]. In addition, myo-inositol in tissues can be depleted by elevated glucose levels, and its dietary supplementation has many benefits including restoring glucose metabolism and improving IS [59, 60], supporting our result. Regarding the reduction of plasma choline in mIR patients, this is the first time that it has been described for IR, to our knowledge. Choline is an essential nutrient for our body, and many studies demonstrated that its metabolism plays an important role in the development of IR and T2DM. In fact, a recent study showed that higher intake of choline is associated with a lower risk of T2DM [61] as well as dietary choline deprivation could result in worsen cardiomyopathy in diabetic rats [62]. All these results supported our previous findings since patients with mIR have a higher IR and cardiovascular risk [8]. Regarding glycerol, it can be used as a cardiac energy substrate [63]. Moreover, several publications associated lower serum creatinine with an increased risk of T2DM [63, 64], in agreement with our study. Furthermore, lower concentrations of plasma creatinine in mIR patients may suggest increased muscle IR, skeletal or striated, and then worsen myocardial conditions according to previous studies [64]. About threonine, a metabolite involved in gluconeogenesis, protein synthesis, and the citric acid cycle for ATP generation, it has a logical reduction in patients with T2DM and even more so if they have the mIR phenotype, since all the metabolic pathways in which it participates are more dysregulated. Moreover, the plasma metabolite threonine is reduced in overweight/obese and T2DMpatients [65]. Finally, plasma VLDL(CH_3_) was the only metabolite increased in mIR patients and it is in accordance with previous trials showing higher concentration in T2DM [66]. In presence of T2DM, increased plasma concentration of VLDL has been related to IR due to an increased VLDL production related to the increase of lipogenesis and reduced ApoB degradation [67]. Therefore, our results are coherent with the knowledge of this metabolite in T2DM.

### Association of GM and plasma and fecal metabolites with myocardial and systemic IR without myocardial phenotyping

After assessing the specific changes in GM, fecal and plasma metabolic profiles between myocardial phenotypes, spearman’s correlation analyses were conducted using all patients to learn more in detail about the relationship of these alterations, as well as to deepen the understanding of myocardial IR disorder. Interestingly, positive relationships were observed between myocardial IS and systemic index of IS, ISI_HEC_, with the genus of sulfate-reducing bacteria *Desulfosporosinus* (Figure 5). This is important because it confirms the deleterious effect of losing this genus in the GM for the evolution of the pathophysiology associated to myocardial IR or systemic IR in T2DM, probably due to its important roles in reducing sulfate to H_2_S, which has received much attention as gaseous modulator of host physiology and mediate increased IS in diabetes [68]. Concerning fecal metabolites, β-galactose was also similarly positively associated with myocardial IS and IS_HEC_. This relationship is very important since this metabolite can be used as a substrate of ATP production [69], and may be preferentially utilized by the myocardium and other tissues to produce energy. Moreover, this metabolite is also present in plasma and has shown a similar positive association with myocardial IS, confirming the importance of β-galactose in the evolution of myocardial IR and its utility as biomarker. Other interesting positive association found were plasmatic VLDL(CH_3_) levels with HOMA-IR and myocardial IR. This association confirmed the well knowledge about the relationship between VLDL and IR [67, 70].

On the other hand, some relationships were obtained exclusively for myocardial IS, including *Coprobacillus cateniformis*, fecal metabolites arabinose, ribose, aspartate and histidine, and the plasma metabolite choline. To our knowledge, this is the first study in patients with T2DM focused on this topic due to the difficulty to determine myocardial IS. Like β-galactose, ribose and arabinose are monosaccharides, and this type of compounds seems to be implicated in the evolution of myocardial IR, despite β-galactose seems to have a more important role. Therefore, this type of monosaccharides needs to be managed in T2DM patients, at least in nutrition or in other products that increase the fecal concentration of this type of monosaccharides. For example, a recent study showed that arabinose treatment could improve IS and reduce HOMA-IR index in obese mice [56] or another proposed ribose to improve metabolism and the cardiac function in T2DM [55]. Therefore, our studies confirm the importance of these sugars in T2DM and need to be studied further to know the underlying mechanisms. Concerning aspartate and histidine, they have also been confirmed to be implicated in IR, and their supplementation in patients may improve the pathophysiology related to IR [53, 54]. Finally, choline is an essential nutrient that could be used to produce energy in the body. It is well-known that in situation of stress, the myocardium could use it as alternative fuel. Therefore, this positive association with myocardial IS could be expected. Moreover, previous studies reinforce the current trial and additionally choline has even been linked to IS in patients [71]. These specific metabolites have potential to be biomarkers, especially β-galactose, for monitoring myocardial IR in T2DM. Despite this, validation in large-scale multi-center cohorts is required.

To know whether the changes of these metabolites were linked with some gut bacteria, relationships between altered gut bacteria and fecal and plasma metabolites were performed these correlations. In all subjects, only one association could be obtained, between *Desulfuromusa* and fecal histidine. This genus of bacteria synthesizes histidine only to meet its own cellular needs and has no specific role in histidine production. However, *Desulfuromusa* degrades organic compounds and reduces sulfur. This process contributes to nutrient recycling, which benefits other microorganisms that can produce histidine and this may be a possible mechanism to understand this association.

### Association of GM and plasma and fecal metabolites with myocardial and systemic IR in mIS and mIR phenotypes

To determine the relationships with each myocardial phenotype, instead of treating the entire population as a whole, we conducted correlation analyses considering only each myocardial phenotype (Figure 5B and 5C).

First, in the population with the mIS or mIR phenotype, unlike the results described above, we did not find a significant correlation between myocardial IS and GM or plasma metabolites. Only the fecal metabolites β-galactose and arabinose maintained consistent associations with myocardial IS in mIS patients. These findings suggest that these metabolites may play an important role in the progression toward myocardial IR in mIS individuals, as such relationships were absent in the mIR phenotype. Notably, we highlight the potential role of β-galactose in the natural history of IR development in T2DM. However, this is the first study to report associations between these fecal monosaccharides and myocardial IS; therefore, further mechanistic investigations and validation studies are required to elucidate their biological roles and assess their potential as biomarkers.

Regarding GM, we did not find any significant associations specifically with myocardial IS. However, we would like to highlight the sulfate-reducing bacterium *Desulfovibrio* had an association with systemic IS_HEC_, and this association is performed in an opposite way in each phenotype, showing its importance in the insulin-sensitization of the myocardium in T2DM. Finally, as expected, plasma VLDL(CH_3_) still had positive correlations with HOMA-IR in mIR patients and coherent with previous findings in systemic IR [67, 70]. It emphasizes the need to pay more attention to plasma VLDL levels in patients with mIR to control the deleterious evolution of T2DM.

Concerning the relationships between altered gut bacteria and fecal and plasma metabolites, we found a negative relationship between *Desulfosporosinus* and plasma β-galactose, and *Thermicanus* and plasma glycerol in mIS patients. *Desulfosporosinus* mainly consumes fermentation products such as lactate that is a precursor of β-galactose, a sugar typically derived from the breakdown of lactose or complex carbohydrates [72]. Therefore, this relationship could come from the degradation of nutrients from *Desulfosporosinus*. This is very important since we have shown that β-galactose plays an important role in myocardial IS. Therefore, this genus of bacteria must be taken into account in the evolution of the pathological history of T2DM in terms of myocardial IR before patients achieving the most pathological phenotype, mIR.

The present study reports several novel and clinically relevant findings that may improve the understanding and management of myocardial IR in T2DM. Nevertheless, the limitations of this study should be acknowledged. First, it lacked a healthy control group, as ethical constraints preclude performing myocardial IR assessments using ¹⁸F-FDG PET/CT in healthy individuals, particularly when repeated measurements are required. Second, the number of patients in each group was limited but appropriate for a proof-of-concept clinical study. Importantly, the groups were well matched in terms of sex, BMI, waist/hip ratio, diet, HbA1c, age, and glucose levels, thereby providing a solid foundation for future investigations.

## 5. Conclusions

This study provides new insights into myocardial insulin resistance in patients with T2DM by revealing alterations in the gut microbiota and metabolome. More specifically, our results show that reduced abundance of *Desulfosporosinus* and *Coprobacillus cateniformis* is associated with myocardial insulin resistance. Moreover, myocardial IR was associated with high levels of a type of VLDL in the plasma and low levels of β-galactose in feces and plasma. Taking into account the relationship between all these variables, we confirm that *Desulfosporosinus* and β-galactose are the key biomarkers for managing myocardial IR in T2DM patients and potential targets for personalized treatments.

## Author Contributions

J.R.H. and M.P.-S. conceived the study and designed the experiments. X.X. performed the majority of experiments, data analysis, and wrote the manuscript, A.C. collected clinical samples and C.A. conducted the blood biochemical analysis. The clinical trial was conducted in the hospital by A.C. and R.S., and the project was administrated by R.S. and J.R.H.. And J.R.H. and M.P.-S. revised the manuscript. All authors have read and agreed to the published version of the manuscript.

## Ethics approval statement

The clinical trial has been registered on https://clinicaltrials.gov/ with the number NCT02248311. This study was conducted according to the tenets of the Helsinki Declaration after approval by the Ethic Committee of the Vall d’Hebron University Hospital (protocol number PR(AG)01/2017). Written informed consent of all participants was collected before the study.

## Acknowledgments

The authors are very grateful to Dr. Julia Baguña Torres, Dr. Daniel García-Leon, and Dr. Bruno Paun for their work in PET/CT image analysis and for providing relevant normalized data. Technician Roso Mares spent a lot of time on the transfer of blood and stool samples, as well as the processing of blood samples, and we are also very grateful to her. At the same time, we thank all the patients involved in the study, as well as the nurses and technicians in the Department of Endocrinology, Department of Biochemistry, Department of Nuclear Medicine and Radiology at Vall d’Hebron University Hospital, for their assistance during the clinical trial.

## Data availability statement

16S rRNA gene sequencing data have been deposited into the NCBI Sequence Read Archive (SRA) database with the accession number PRJNA810955 (https://www.ncbi.nlm.nih.gov/sra).

